# metabaR : an R package for the evaluation and improvement of DNA metabarcoding data quality

**DOI:** 10.1101/2020.08.28.271817

**Authors:** Lucie Zinger, Clément Lionnet, Anne-Sophie Benoiston, Julian Donald, Céline Mercier, Frédéric Boyer

## Abstract

1. DNA metabarcoding is becoming the tool of choice for biodiversity studies across taxa and large-scale environmental gradients. Yet, the artefacts present in metabarcoding datasets often preclude a proper interpretation of ecological patterns. Bioinformatic pipelines removing experimental noise have been designed to address this issue. However, these often only partially target produced artefacts, or are marker specific. In addition, assessments of data curation quality and the appropriateness of filtering thresholds are seldom available in existing pipelines, partly due to the lack of appropriate visualisation tools.
2. Here, we present **metabaR**, an R package that provides a comprehensive suite of tools to effectively curate DNA metabarcoding data after basic bioinformatic analyses. In particular, **metabaR** uses experimental negative or positive controls to identify different types of artefactual sequences, i.e. reagent contaminants and tag-jumps. It also flags potentially dysfunctional PCRs based on PCR replicate similarities when those are available. Finally, **metabaR** provides tools to visualise DNA metabarcoding data characteristics in their experimental context as well as their distribution, and facilitate assessment of the appropriateness of data curation filtering thresholds.
3. **metabaR** is applicable to any DNA metabarcoding experimental design but is most powerful when the design includes experimental controls and replicates. More generally, the simplicity and flexibility of the package makes it applicable any DNA marker, and data generated with any sequencing platform, and pre-analysed with any bioinformatic pipeline. Its outputs are easily usable for downstream analyses with any ecological R package.
4. **metabaR** complements existing bioinformatics pipelines by providing scientists with a variety of functions with customisable methods that will allow the user to effectively clean DNA metabarcoding data and avoid serious misinterpretations. It thus offers a promising platform for automatised data quality assessments of DNA metabarcoding data for environmental research and biomonitoring.

## 1 Introduction

DNA metabarcoding coupled with high-throughput sequencing is currently revolutionising the way we describe biodiversity across environments and taxa, and is therefore becoming a tool of choice for basic and applied research, as well as for biomonitoring applications (Deiner et al., 2017; Taberlet et al., 2018; Cordier et al., 2020). In recent years, various bioinformatic pipelines and tools have been developed to handle DNA metabarcoding data. These include e.g. **qiime** (Caporaso et al., 2010; Estaki et al., 2020), **OBITools** (Boyer et al., 2016; Taberlet et al., 2018), **vsearch** (Rognes et al., 2016), or **dada2** (Callahan et al., 2016). These bioinformatic packages typically perform bioinformatic analyses such as sequence alignment, clustering into MOTU (Molecular Operational Taxonomic Unit), noise data removal, or taxonomic assignment and ultimately produce a MOTU by sample matrix. This matrix, similar to the community table of community ecologists, can then be used to reveal patterns of alpha and beta diversity with more classical ecological R packages such as **vegan** (Oksanen et al., 2019) or **adiv** (Pavoine, in press), or with packages dedicated to microbiome analyses (e.g. **phyloseq**, McMurdie & Holmes, 2013).

While the aforementioned bioinformatic tools have been heavily used for analysing the ever-expanding number of DNA metabarcoding data, they also present a certain number of limitations. DNA metabarcoding generates numerous experimental biases besides PCR/sequencing errors and chimeras, which range from field or laboratory contaminations through to tag-jumps (Table 1; reviewed in Taberlet et al., 2018; Zinger et al., 2019). The treatment of these artefact is often missing in DNA metabarcoding studies, even though they can substantially affect ecological inferences (Sommeria-Klein et al., 2016; Frøslev et al., 2017; Calderón-Sanou et al., 2019). Such artefacts can only be flagged and corrected by including experimental controls and experimental replicates throughout the data production process. However, existing bioinformatic pipelines only deal with PCR/sequencing errors, and do not make use of experimental controls to filter out potential contaminants or artefacts. Second, these bioinformatic pipelines often lack tools to monitor and evaluate the bioinformatic data filtering process. As a result, it is often difficult to tune data filtering parameters, and users are therefore led to using default settings even when these are suboptimal. Finally, DNA metabarcoding data are in essence multidimensional, as they encompass MOTUs, PCR product, and biological sample information. This multi-fold information, often stored in separate tables, is not easily handled by most R packages for data analyses (but see e.g. **phyloseq**). As such, we currently lack effective tools to transition from bioinformatic pipelines to ecological R packages.

**Table 1.** Overview of DNA metabarcoding experimental artefacts

| Experimental bias | Description |
| --- | --- |
| PCR/sequencing errors | Any MOTU resulting from base misincorporation by DNA polymerase during PCR amplification or sequencing, or base miscalling during sequencing. |
| Contaminants | Any MOTU external to the biological sample, typically from lab reagents. Such contamination can occur at all stages of the data production, i.e. field work, DNA extraction, PCR amplification, and library preparation. |
| Tag-jumps | MOTU of which presence is genuine at the sampling sites, but erroneous in a given sample/PCR product due to a switch of so called "tag" or "library index", i.e. a nucleotidic kmer allowing to reassign the amplicon to the amplicon to its sample/PCR reaction of origin. |
| Degraded sequences | Any sequence or MOTU such as primer dimers, or chimeras from two or multiple parents. Usually largely differ from any known sequence. |
| Dysfunctional PCRs | Any PCR product yielding a low amount of sequences or an irreproducible signal. |

To bridge this gap, we developed **metabaR**, an R package that enables the post-processing and filtering of DNA metabarcoding data already processed through bioinformatic pipelines so as to improve downstream ecological inferences. It is designed to take advantage of negative controls, positive controls and PCR replicates when available to efficiently flag and remove artefactual MOTUs or dysfunctional PCRs. It is implemented in the R statistical programming environment (R Core Team, 2020) which provides flexible analytical tools coupled with powerful graphical capabilities. **metabaR** uses these properties to provide highly customisable functions, as well as effective visualisation of DNA metabarcoding data in their experimental context. Hence, it is of direct use for any practitioner of DNA metabarcoding techniques with basic skills in R programming.

## 2 Data structure, import/export, and manipulation

**metabaR** performs the analysis of DNA metabarcoding data by taking into account its multidimensional nature. The central object of the package is a `metabarlist`, an R list composed of 4 interconnected tables (Fig. 1): (i) `reads`, a table that stores the read abundance of MOTUs in each PCR product, (ii) `motus`, a table which stores any information relative to each MOTU in the dataset (e.g. taxonomic information), (iii) `pcrs`, a table which stores any information relative to each PCR reaction (e.g. if it is a sample or an experimental control, what are the primer used, etc.), and (iv) `samples`, a table which contains any metadata relative to the biological sample from which the PCR reaction was obtained (e.g. geographic coordinates, abiotic parameters, etc.). This object can be generated from outputs of various bioinformatic pipelines such as **vsearch**, **qiime** or **OBITools** through a set of data-import functions. These include two generic functions, `tabfiles_-to_metabarlist` and `biomfiles_to_metabarlist` that import files in csv or BIOM (Biological Observation Matrix) format, and the more specific `obifiles_to_metabarlist` function adapted for **OBITools** outputs. We also provide the `metabarlist_generator` function, which facilitates `metabarlist` building directly from objects in the R environment. The package also provide a tool, `silva_annotator`, which imports **SILVAngs** (Quast et al., 2012) taxonomic output files to complement the `motus` table for more specific applications.

**Figure 1.**
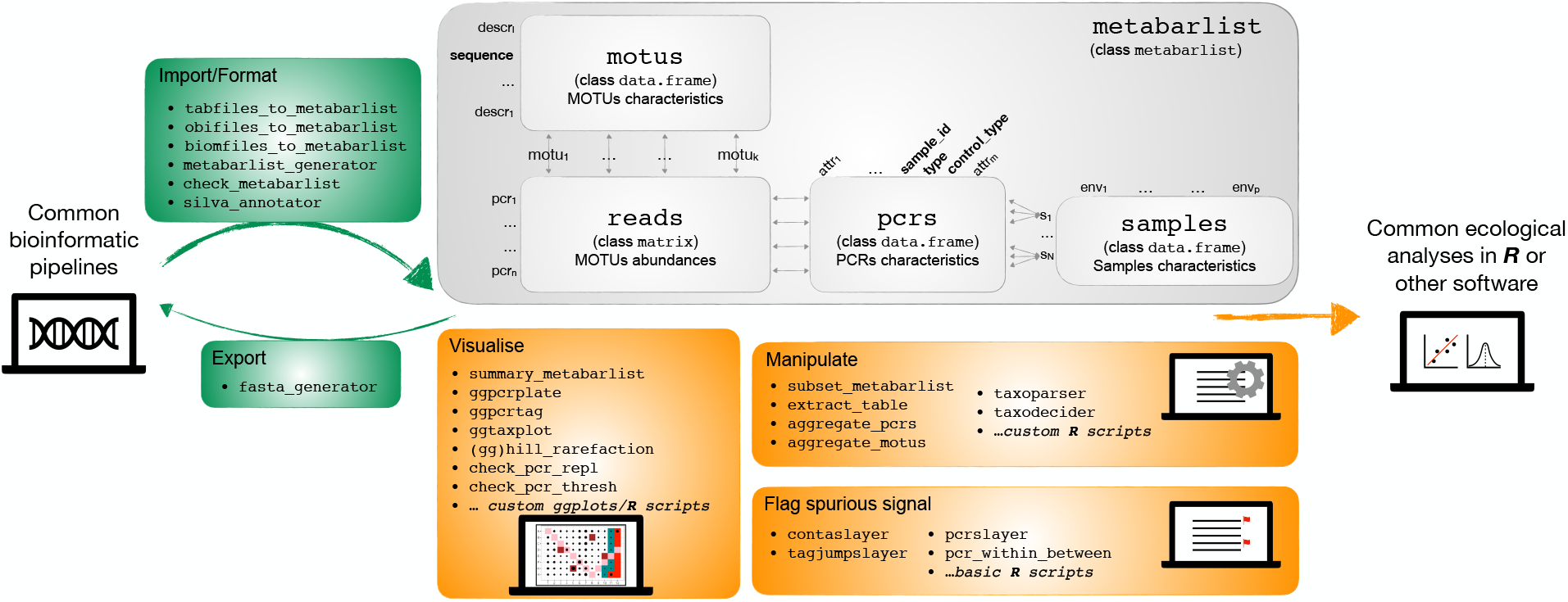
Overview of the **metabaR** package data structure (grey box) and functions (green and orange boxes). Mandatory fields in each table of the metabarlist are indicated in bold. More details are available in the help page of `check_metabarlist`.

All these import tools use the function `check_metabarlist`, which verifies whether the imported or created `metabarlist` fulfills a set of mandatory properties for the package to work. The function returns a warning message with guidance to the user when the format is incorrect.

Any table of the `metabarlist` can be amended easily with R commands non specific to the **metabaR** package. For example, the `reads` matrix can be transformed into relative abundance data with `rowSums` or the `decostand` function of the **vegan** package. Likewise, any column can be added to the data frames `pcrs`, `motus`, or `samples` by using basic R commands.

The `metabarlist` object can be manipulated for different purposes. It can be subsetted with `subset_metabarlist` with customisable criteria relating to any table of the `metabarlist`. The user can also aggregate read counts based on MOTU criteria with `aggregate_motus`, such as for obtaining community data at higher taxonomic ranks than the OTU level. Similarly `aggregate_pcrs` can be used to aggregate read counts based on PCR related criteria, typically to aggregate technical PCR replicates at the sample level.

We also include two functions to facilitate the customisation of taxonomic information. The first, `taxoparser` is a simple tool that parses full or partial taxonomic paths generated during upstream bioinformatic processing. The function `taxodecider` enables users to process taxonomic assignment for the same MOTU from multiple databases. For example, building a custom reference database is often recommended, since including species from the regional species pool increases the reliability of taxonomic assignments (Taberlet et al., 2018). However, these databases are often incomplete and it is common to run in parallel annotation tools with more generalist reference databases such as EMBL (https://www.embl.org/). The`taxodecider` function allows users to merge different annotations based on assignment scores and by giving priority to assignments from the user’s preferred reference database, usually the one for which taxonomic and sequence information is the most reliable.

Finally **metabaR** has different export tools. First, `fasta_generator` exports sequences in the fasta format where the user is free to add any information from the `metabarlist` to the sequence header. This function can be of use when following the implementation of **metabaR** functions, it becomes apparent that specific bioinformatic data curation procedures require retuning. The package does not provide other export functions, since the `metabarlist` is a simple R `list` that can be directly exported with the R base function saveRDS. Alternatively, any table may be extracted from the `metabarlist`with `extract_-table`, before R’s `write.table` function is used for export.

## 3 Example dataset

The package contains a dataset, named `soil_euk`, which is a typical output of a DNA metabarcoding experiment. It is used in the package help and the vignette to illustrate the functions of **metabaR**. `soil_euk` is a `metabarlist` and contains the abundance of 12,647 MOTUs obtained from 384 PCRs, corresponding to a total of 64 biological samples. The dataset also includes different information on MOTUs, PCRs, and samples. The dataset was generated from soil and litter samples collected in two tropical forests in French Guiana, from which a variable region of the 18S rRNA gene was amplified by PCR and sequenced on an Illumina platform. The data was then processed with the **OBITools** bioinformatic pipeline. Detailed information regarding the generation of this dataset is available in the `soil_euk` help page.

## 4 Visualisation

Appropriately representing DNA metabarcoding data visually is a prerequisite to assessing the quality of the data or of the curation process. Such assessments require going beyond representing dataset characteristics such as sample sequencing depth or richness in MOTUs using standard boxplots and histograms. Here, we developed two functions, `ggpcrplate` and `ggpcrtag`, to represent data set characteristics in their experimental context, i.e. the PCR plate. Their input consists of a `metabarlist` and a function pre-encoded in **metabaR** or designed by the user to be applied to the input `metabarlist` so as to enable the plotting of numerous dataset characteristics. Such visualisation can enable the identification of potential experimental problems, such as pipetting or tag/primer issues as exemplified in Fig. 2.

**Figure 2.**
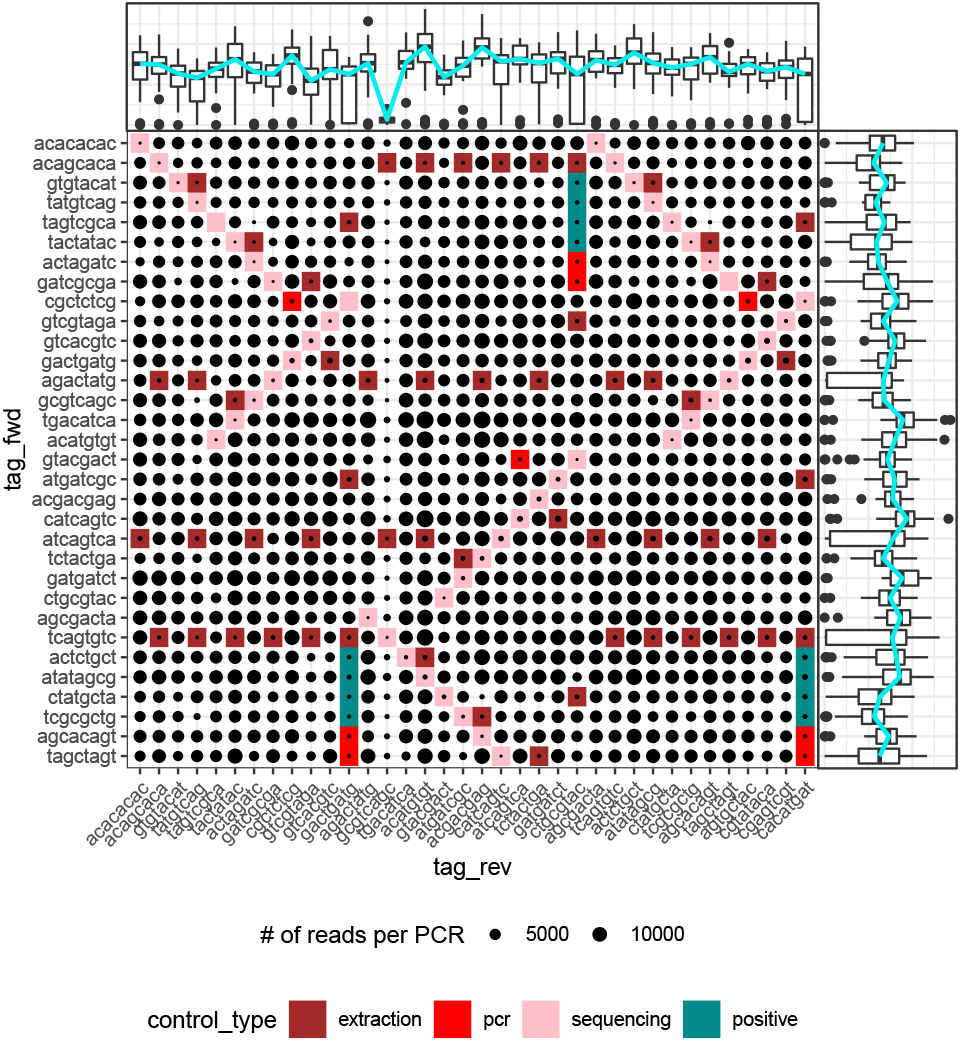
Example of an output from `ggpcrtag` with a problematic DNA metabarcoding dataset exhibiting low amounts of reads for all PCR reactions conducted with the reverse primer including the tag “gcgtcagc”. Upper and right boxplot show the total value of the variable of interest (here number of reads) across all PCRs using a primer with the same tag. The figure also shows what experimental design was used for this particular dataset (controls type and locations in a 4 × 3 PCR plate set up).

The taxonomic composition of DNA metabarcoding data is also often difficult to represent because taxonomic assignments are seldom available at a uniform taxonomic level. This problem usually results from either the incompleteness of reference databases, or as a result of the inherent variation of DNA markers in taxonomic/phylogenetic resolution across lineages. To facilitate the visualisation of the sample or experiment’s community composition in this context, we developed the function `ggtaxplot`, dependant on the **igraph** R package (Csardi & Nepusz, 2006). This function plots taxonomic trees where each node corresponds to a taxon, with node size and colour corresponding to the taxon number of reads and diversity in MOTUs (Fig. 3).

**Figure 3.**
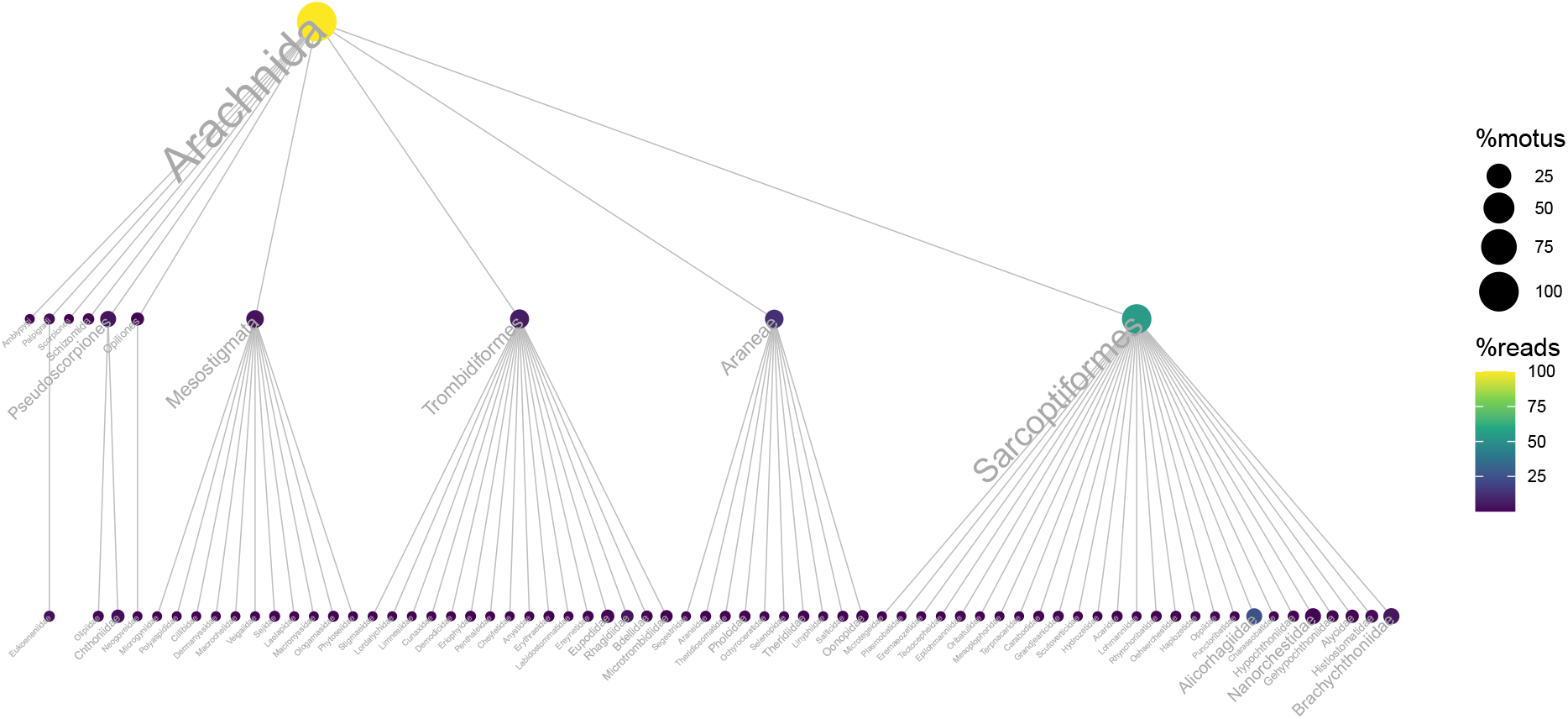
Example of an output from `ggtaxplot` using the `soil_euk` dataset, focusing on Arachnida MOTUs. Each node corresponds to a taxon, node size to the proportion of MOTUs, and node color to the proportion of read counts.

Finally, rarefaction curves are routinely used with DNA metabarcoding data to assess whether the MOTU diversity of each PCR reaction or sample is appropriately covered by sequencing depth. The `hill_rarefation` function and its plotting complement `gghill_- rarefaction` build rarefaction curves using three indices included in the Hill numbers framework (Hill, 1973; Chao et al., 2014), which have been shown to provide good estimates of alpha diversity for DNA metabarcoding data (Alberdi & Gilbert, 2019; Calderón-Sanou et al., 2019). More specifically, these functions estimate MOTU richness, the exponential of the Shannon index, and the inverse of the Simpson index; as well as Good’s coverage index (Good, 1953) at different sequencing depths chosen by the user.

All visualisation tools used in **metabaR** are based on `ggplot2` and **cowplot** R packages (Wickham, 2016; Wilke, 2019) for greater flexibility.

## 4 Data curation tools

Numerous bioinformatic tools allow the curation of DNA metabarcoding data to account for PCR and sequencing errors. By contrast, only a few (e.g. **LULU**, (Frøslev et al., 2017)) deal with other types of artefactual MOTUs (Table 1). **metabaR** includes three functions which each target a particular type of noise data. To allow users to evaluate the downstream impacts of removing identified noise data, two of these only flag potential spurious objects in the output rather than removing them directly.

The `tagjumpslayer` function targets artefacts called “tag-jumps”, “tag-switches” or “cross-talks” (Table 1, Schnell et al., 2015; Esling et al., 2015; Edgar, 2017), which generate a noise similar to cross-sample contaminations but at the scale of the whole sequencing library, hence homogenising the data. The `tagjumpslayer` function aims to reduce this noise by removing a MOTU in a given PCR product when its relative abundance over the entire dataset is below a given threshold. This threshold can be empirically chosen by testing the effect of varying curation thresholds on the MOTU and read counts in the dataset in general and, when available, in the sequencing negative controls (i.e. unused tag or library index-combinations) in particular.

The effect of these tag-jumps can complicate the detection of external contaminants, such as those occurring in laboratory reagents (Salter et al., 2014). An approach which only consists in the detection of MOTUs present in experimental negative controls would ignore tag-jumps, and can result in the removal of the most abundant genuine MOTUs from the dataset. However, in negative controls, contaminants should be preferentially amplified in the absence of competing DNA, which is unlikely to be the case in biological samples. The `contaslayer` function relies on this assumption and detects MOTUs whose relative abundance across the whole dataset is highest in negative controls.

Finally, the `pcrslayer` function aims to identify potential failed PCR reactions by comparing the dissimilarities in MOTU composition within a biological sample (i.e. between PCR replicates, hereafter *dw*) vs. between biological samples (hereafter *db*). It relies on the assumption that PCR replicates from a same biological sample should be more similar than two different biological samples (*dw < db*). A PCR replicate having *dw* above a given dissimilarity threshold, defined automatically by the function based on the distribution of *dw* and *db*, is considered to be an outlier. The function can be run with any dissimilarity index. Several functions are provided along with `pcrslayer`, such as `check_pcr_repl`, which draw an ordination of PCR replicates; as well as `pcr_within_between` and `check_pcr_thresh` which compute and represent the distribution of *dw* and *db*.

In addition to the identification and flagging of artefacts provided by these functions, other issues such as PCRs with shallow sequencing depths, MOTUs that are not targeted by the primers or those with too low taxonomic assignment scores, can also be flagged with R base functions (detailed in the package accompanying vignette).

## 5 Conclusions

The **metabaR** package provides a much needed tool at the interface between bioinformatic pipelines and ecological analyses to evaluate the quality of data and of the curation process, prior to offering further curation of commonly overlooked artefacts. We also provide a vignette along the package that constitutes for new users a good starting point to build their own quality assessment and filtering of DNA metabarcoding data: it highlights all the recommended steps and possible uses of experimental controls to clean the data. The **metabaR** package and its vignette will contribute in improving data quality standards in the field, ease the analysis of DNA metabarcoding data and will therefore help to broaden the use of eDNA-based analyses of biodiversity.

## Acknowledgements

We are deeply indebted to Eric Coissac for stimulating discussions that led to the development of this package, and are also grateful to Jerôme Chave and Wilfried Thuiller for supporting this work. We thank Pierre Taberlet and Heidy Schimann for providing data, as well as Irene Calderón-Sanou, Camille Martinez-Almoyna, Jerôme Murienne and Renato A. Ferreira de Lima for practical discussions on - and/or testing of - earlier versions of the package. We also thank Chris Bowler for providing informatics equipment to ASB. The work was funded by the METABAR (ANR-11-BSV7-0020) and GlobNets (ANR-16CE02-0009) projects, and has benefitted from “Investissement d’Avenir” grants managed by Agence Nationale de la Recherche (CEBA: ANR-10-LABX-25-01; TULIP: ANR-10-LABX-0041; OSUG@2020: ANR-10-LABX-56).

## Authors contribution

LZ, FB, and CL conceived and wrote the package. ASB and CM contributed to the writing of functions and ASB and JD to the writing of the documentation and vignette. LZ wrote the manuscript with inputs from all co-authors.

## Data availability statement

The **metabaR** package is available on GitHub at https://github.com/metabaRfactory/metabaR. We also provide a full description of the package functions, as well as a step by step tutorial (R vignette) describing the package basic use at https://metabarfactory.github.io/metabaR. The example dataset is provided within the package in .rds, .biom, and .txt formats.

